# Stochastic model of contact inhibition and the proliferation of melanoma *in situ*

**DOI:** 10.1101/110007

**Authors:** Mauro César Cafundó Morais, Izabella Stuhl, Alan U Sabino, Willian W Lautenschlager, Alexandre S Queiroga, Tharcisio Citrangulo Tortelli, Roger Chammas, Yuri Suhov, Alexandre F Ramos

## Abstract

Contact inhibition is a central feature orchestrating cell proliferation in culture experiments; its loss is associated with malignant transformation and tumorigenesis. We performed a co-culture experiment with human metastatic melanoma cell line (SKMEL-147) and immortalized keratinocyte cells (HaCaT). After 8 days a spatial pattern was detected, characterized by the formation of clusters of melanoma cells surrounded by keratinocytes constraining their proliferation. In addition, we observed that the proportion of melanoma cells within the total population has increased. To explain our results we propose a spatial stochastic model (following a philosophy of the Widom-Rowlinson model from Statistical Physics and Molecular Chemistry) which considers cell proliferation, death, migration, and cell-to-cell interaction through contact inhibition. Our numerical simulations demonstrate that loss of contact inhibition is a sufficient mechanism, appropriate for an explanation of the increase in the proportion of tumor cells and generation of spatial patterns established in the conducted experiments.

## Introduction

Despite the accumulated knowledge of experimental results on contact inhibition as an *in vitro* manifestation of homeostatic cell density control in normal tissues, the use of quantitative tools to understand its role in the growth of cancer *in situ* is only in its infancy ^1,2^. Contact inhibition can be described as the decrease of proliferation rates when the cell density increases. At the molecular level, intercellular adhesion mediated by E-cadherin (CDH1) serves as negative regulator of the cell proliferation signal by recruiting *β*-catenin to adherens junctions and thus repressing the transcriptional activation of proliferative genes ^3^. Another mechanism is manifested by the interplay between the cyclin-dependent kinase inhibitors p27 and p16, tumor suppressor proteins. In a recent experiment, it has been demonstrated that p16 is responsible for a stronger sensitivity to contact inhibition in combination with p27^4^. This has provided a clue about the cancer resistance of naked mole-rats as compared to human and murine fibroblasts. While in human and murine fibroblasts, the p16 expression is attenuated when cell density increases, the coordinated expression of both p16 and p27 in naked mole-rat fibroblasts rendered these cells more resistant to malignant transformations. The naked mole-rat fibroblast growth is strongly inhibited by cell-to-cell contacts, while the growth of human and murine fibroblasts is supported even at higher densities.

In this study we suggest that cells whose growth overcome contact inhbition, i.e. cells tolerating higher cellular densities, are more resistant to allelopathic effects of their neighbours, and can therefore be called *allelophylic* (*allelo*, the other; *phylia*, affinity). This neccesitates the proposition of a theoretical model helping to account for different degrees of contact inhibition and quantify their role in the carcinogenesis. Mathematical models have been useful for the analysis of high throughput data^5–7^, identification of tumorigenesis^8–12^, angiogenesis^13^, cancer invasiveness ^14–16^, therapy design^17–21^, and for the detection of the effects of intrinsic randomness of cellular phenomena in cancer^11,22^. In particular, new theoretical approaches have been developed to show that contact inhibition occurs through both cell-to-cell contacts and mechanical constraints reducing cells to the substrate adhesion area ^23^. In this paper we present theoretical and experimental frameworks designed to investigate the modulated contact inhibition and its role in the formation of a carcinoma *in situ* and to demonstrate that allelophilic properties of cancer cells is a key feature for their uncontrolled proliferation.

## Results

### Keratinocytes and melanoma cells co-culture proliferation

To evaluate the cell proliferation, the human metastatic melanoma (SK-MEL-147) and human immortalized keratinocytes (HaCaT) cell lines were selected for co-culture experiments. The choice of these cells allows us to mimic the interaction between the skin basal layer cells and the melanoma. Another reason for selecting these cell lines was to compare the co-culture development with patterns produced through a stochastic model dynamics. The latter involves a cell line that shows a distinctive degree of contact inhibition (a property of HaCaT) and another cell line that is highly tolerant, i.e., displays a loss of contact inhibition (which is a characteristic of SK-MEL-147). In the supplementary material we collect results of our experiments. At the post confluence stage the carrying capacity of HaCaT is at 1779.56 ± 130.47 cells/mm^2^ while for SK-MEL-147 it equals 5043.51 ± 316.47 cells/mm^2^ (see *Data Fitting* section and Fig. S1 at the Supplementary Information file). This demonstrates higher density levels achieved by melanoma cells (Fig. 1) confirming their distinctively lower degree of contact inhibition in comparison with keratinocytes. A similar phenomenon was observed in a different situation in^4^.

**Figure 1.**
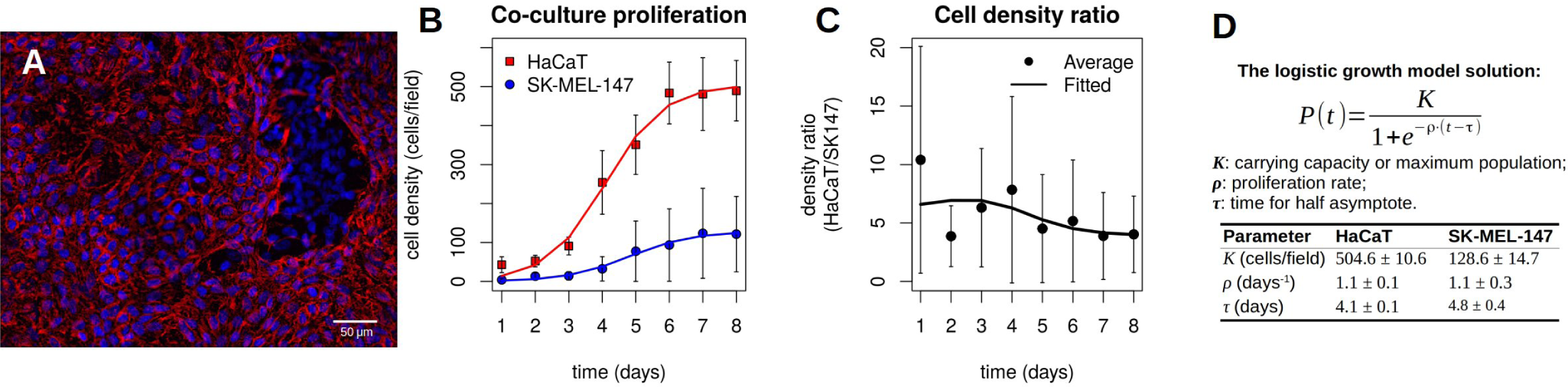
HaCaT and SK-MEL-147 cells co-culture proliferation. **A**. Immunofluorescent staining of E-cadherin (CDH1) on HaCaT and SK-MEL-147 co-culture. Cells were fixed and stained with the mouse anti-CDH1 (red). The secondary antibody was the goat anti-mouse Alexa Fluor 546, and nuclei were stained with Hoechst 33258 (blue). The difference in the CDH1 expression presented by SK-MEL-147 was used to distinguish between the two cell lines in co-culture images. When confluence was reached, after 4 days, it was possible to observe SK-MEL-147 domains surrounded by HaCaT cell layers. **B**. The cell proliferation curves of HaCaT and SK-MEL-147 cells in the co-culture. Cells were counted in 30 random fields of view every day. Blue circles indicate SK-MEL-147 while red squares indicate HaCaT averages of cells/field. Error bars correspond to the standard deviation. Solid lines indicate fitted data from the logistic growth model. **C**. The cell density ratio (HaCaT:SK-MEL-147). The experiments started with a cell density proportion of 10:1 which decreased to ~ 4:1, despite maintaining the same proliferation rates. **D**. The solution for the logistic growth model and parameter value estimates. The data were fitted by using the nls() function from R software.

At the initial stage of the co-culture experiments, cells were seeded at 250 cells/mm^2^, at a proportion of keratinocytes to melanoma of 10:1, in a monolayer on a 24-well plate dish with coverslips. The co-culture was allowed to proliferate for eight days. The monolayer structure enabled us to investigate the role of contact inhibition in the cell proliferation at a quantitative level. After four days in the co-culture, cells reached confluence, and it was possible to observe the formation of growing melanoma clusters. These clusters are constrained by layers of keratinocytes cells, of density somewhat higher than normal (Fig. 1**A**). To evaluate the cell population growth, we counted the number of cells in images from 30 locations on the plate for each day of experiment. The obtained data were fitted by using the logistic growth model (Fig. 1**B**). The parameter *ρ* indicates the cell population growth rate, the maximum population density is denoted by *K*, and the time when the cell density equals *K*/2 is τ. The values for the fitting for each cell line are shown in Fig. 1**D**. Note that *ρ* and τ can be made (approximately) the same for both cell lines, while the ratio between the maximum densities is ~ 4. The change in time of the ratio between the two cell population densities is shown in Fig. 1**C**. One may also note that the proportion of HaCaT cells density decays from ~ 10:1 in the first-day measurement to ~ 4:1 at the eighth day. Although the logistic model gives a proper fit for the experimental data, it yields no insight on the mechanisms underpinning the increase on the proportion of melanoma cells. To this end we propose a more detailed model aiming at providing a qualitative description of the observed data.

#### A stochastic model of cell proliferation for a cell-to-cell interaction through the contact inhibition

A mathematical model explaining the above experimental results is related to the Widom-Rowlinson model^24–26^. Since the data refer to a monolayered co-culture arrangement, we can focus on a two-dimensional system composed of two cell types interacting as colored billiard balls of different sizes, where each cell type has two *exclusion areas* surrounding it (one for balls of the same color, the other for balls of different colors). Here type 1 are tumor/cancer cells (SK-MEL-147) and type 2 are healthy/normal cells (HaCaT). The cells are placed at the vertices of a two-dimensional grid Λ formed by unit squares. The distance between two vertices is represented by the minimal number of unit edges connecting them (a graph metric). An admissible configuration of this system satisfies the following rules: (i) each vertex is occupied by at most one cell; (ii) the distance between two cells *i* and *j*, with *i*, *j* = 1, 2, is a positive integer not less than a given value *D*(*i*, *j*), referred to as exclusion distance/diameter between cell types *i* and *j*. In fact, *D*(*i*, *j*) represents the minimal allowed distance between sites occupied by cells of types *i* and *j*.

A formal mathematical description of the model and related results are contained in the supplementary information (see Theorem 1).

The experiments show a close relationship between the degree of contact inhibition among cells and the cell density. It indicates a possible biological interpretation of self-exclusion diameters *D*(*i*, *j*): they measure the degree of contact inhibition of melanoma and keratinocytes, respectively. A greater value of the exclusion diameter *D*(*i*, *i*) indicates a higher degree of contact inhibition. Intuitively, it can be understood in terms of dense-packing the grid Λ by a single cell type. As the self-exclusion diameter of the cell increases, the maximal proportion of occupied vertices reduces. In other words, the cell density within the grid is inversely proportional to *D*(*i*, *i*).

For the simulations, the minimal allowed distances were chosen to be *D*(1, 1) = 1, *D*(1, 2) = *D*(2, 1) = 3, and *D*(2, 2) = 2; this indicates that the smallest exclusion occurs between two tumor cells. Pictorially, a pair of healthy cells are twice less allelophilic and a healthy-tumor pair is three times less allelophilic than the pair of tumor cells. A diagram presenting features of an admissible configuration is shown on Fig. 2**A**: here normal (tumor) cells are represented as red (blue) colored circles. Note that our choice of *D*(1, 1) allows the tumor cells to occupy nearest-neighbor vertices. The Fig. 2**B** shows a magnification of an admissible configuration on a 25 × 35 grid as obtained in the course of a simulation. The spatial pattern is characterized by the formation of clusters of tumor cells surrounded by a sea of normal cells separated by a layer of empty vertices. Such a pattern resembles the keratinocytes and melanoma spatial arrangement observed experimentally, as shown in Fig. 1**A**. As was stated, the exclusion diameter *D*(*i*, *j*) is interpreted as a degree of contact inhibition between cell types *i* and *j*: the smaller *D*(*i*, *j*) is the lower the degree of contact inhibition.

**Figure 2.**
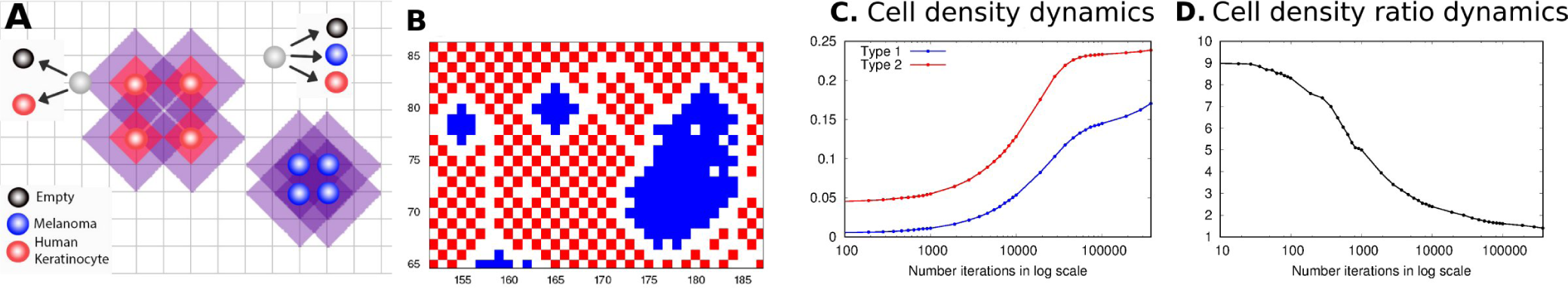
An admissible configuration and the density of the stochastic model. **A**. Two possible state changes of the Markov chain for the minimal distances *D*(1,1) = 1, *D*(1, 2) = *D*(2,1) = 3, *D*(2,2) = 2. The red/blue circles indicate the normal/tumor cells placed at sites/vertices of a square grid. The four blue circles positioned at neighboring vertices represent a part of a densely-packed configuration for tumor cells. The four red circles surrounded by diagonal red squares represent a part of a checker-board densely-packed configuration for normal cells: their inhibition level prevents any further concentration. Each diagonal red square around a red circle marks an exclusion area for other red (healthy) cells. The larger purple region around the red circles covers the vertices forbidden for blue (tumor) cells in the vicinity of healthy ones. The gray circles mark two randomly selected empty vertices, located in a shadowed and non-shadowed region, respectively. The arrows from circles 1 and 2 point to possible states at the next transition: (i) vertex 1 may remain empty or become occupied by a red cell (but not blue), since it is located in a purple shadowed area; (ii) vertex 2 is in a non-shadowed area and may remain empty or become occupied by either a red or blue cell. **B**. A simulated configuration: blue (red) dots indicate tumor (normal) cells. After a sufficiently long time, a spatial pattern is formed: normal cells surround tumor cell clusters, with empty separating layers. Note similarities with experimental images in Fig. 1**A**. Simulations were done on a 200 × 200 grid with empty borders. The division rates are *α*_1_ = *α*_2_ = 0.1, the degradation rates *ρ*_1_ = *ρ*_2_ = 0.01, the migration rates *δ*_1_ = *δ*_2_ = 0.001. **C**,**D**. The number of time steps of the Markov chain: **C** presents the dynamics of the densities for cell types 1 and 2, in colors blue and red; **D** shows the ratio of the densities.

##### Stochastic dynamics of cell proliferation: a Markov chain on admissible configurations

We consider three possible transitions for the *i*-th cell type: division, migration, and death, occurring at a rate *α_i_*, *δ_i_*, and *ρ_i_*, respectively, with *i* = 1, 2. For simplicity, quiescence of cell type *i* is considered to happen at rate *α_i_* but a general version of the model might take different values for that rate. Cell transformations from normal to tumor is neglected due to the time scale of the experiments, and because cell lines used in experiments are from different embryonic origins. We treat the randomness of cell proliferation in terms of a discrete-time Markov chain on admissible configurations on a square grid of size *L* × *L* where *L* = 200. A given vertex of the grid can assume a state 0, 1, or 2 indicating that it is empty (0), occupied by a tumor cell (1), or by a normal cell (2).

The transition from a current state towards the next one begins with selecting a vertex *x* ∈ A with probability *L*^−2^ and verifying its state as occupied or empty. (i) If *occupied* by a cell type *i*, vertex *x* may remain occupied (quiescence) by it with probability *α_i_*/ *Q* or becomes empty with probability 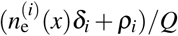. Here 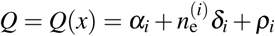 and 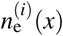 is the number of empty vertices at distance *D*(*i*, *i*) from *x*, which can receive the migrated cell in accordance with the admissibility rule. In case of migration (occurring with probability 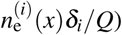, the vertex receiving the moving cell from *x* is chosen with probability 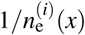, and it becomes occupied by the migrating cell. (ii) If vertex *x* is *empty*, it remains empty with probability (*ρ*_1_ + *ρ*_2_)/*R*, and notice that this probability might be defined in terms of different rates and this choice was done aiming at avoiding introduction of more parameters. Next, a cell type *i* may occupy *x* with probability 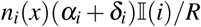 due to a cell division (occurring with probability 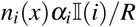) or migration (having a probability 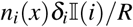). Here 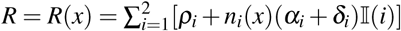, and *n_i_*(*x*) stands for the number of vertices currently occupied by cell type *i* among the nearest allowed neighbors of *x*. The migrating cell is chosen with probability 1/*n_i_*(*x*) and its vertex becomes empty after migration. Further, 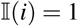 or 0 when vertex *x* can or cannot be occupied by a cell of type *i* without violating the admissibility condition. If *R* = *ρ*_1_ + *ρ*_2_, i.e. vertex *x* cannot be occupied by any type of cell, it remains empty.

The proposed structure of Markov chain transitions gives a rather coarse-grained (and simplified) approximation to possible dynamics of the co-culture cell population. In particular, the outcomes of cell division take into account geometric characteristics of the current cell configuration only partially. (Through numbers 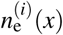 and *n_i_* (*x*).) It is not clear to what extent these simplifications change the overall picture of the co-culture evolution; we consider them as an initial step in the study of mathematical models of such an evolution. Nevertheless, even such a simplified model shows a similarity between curves in Figs. 1 and Fig 2.

##### In the stochastic dynamics, tumor cells increase their proportion due a low degree of contact inhibition

The results of simulations based on the stochastic dynamics are analyzed in terms of the *i*-th cell type density, that is, the proportion of vertices of the grid which are occupied by the *i*-th cell type. Fig. 2C shows the dynamics of the normal and the tumor cell populations. The cell densities are shown on the vertical axis and the time steps of the Markov chain are marked in a logarithmic scale on the horizontal axis. Here one should not confuse the Markov time steps with experimental time scales. In the co-culture experiment there are multiple cell state transitions going on in parallel while in the simulation we take at most one transition per a single time step. An initial configuration is chosen with a the type 1 (tumor) cell density 0.01, type 2 (normal) cell density 0.09 and the total cell population density 0.1. The transition rates are made the same for both cell types, and only the self-exclusion diameters *D*(1, 1) = 1 and *D*(2, 2) = 2 are different, with normal cells having the self-exclusion diameter twice as long as tumor cells. (The choice of the cross-exclusion diameter *D*(1, 2) = *D*(2, 1) = 3 seems to play a lesser role here.) We avoid border effects by maintaining them empty during an entire simulation. The red (or blue) lines dots indicate the normal (or tumor) cell population density. During the time interval of the simulations there is a variation in the ratio of normal to tumor cell-densities, as presented on Fig. 2D. Since all proliferation rates of both cell types are the same, such a change should be attributable to the difference on the values of their exclusion diameters. In biological terms, it means that a low degree of contact inhibition favors the tumor cells population to increase its proportion within the total cell population.

##### Mutual contact inhibition promotes the formation of cell clusters

Fig. 3 shows six frames of simulated admissible configurations of the Markov chain on the grid. An initial configuration is presented on frame **A** while a steady state configuration pattern is shown on frame **F**. The intermediate frames, **B**, **C**, **D**, and **E**, demonstrate transient admissible configurations. Frames **B** and **C** display an initial formation of small clusters of tumor cells. The configuration at the time step 10^5^ is shown on frame **D**; it corresponds to the cell population densities observed in Fig. 1**A**. The magnification of this configuration at a given region of the domain (seen in Fig. 2**B**) resembles the picture in Fig. 1**A**: one may observe multiple small clusters of tumor cells surrounded by a sea of normal cells. This spatial pattern is reverted in the course of time as shown on frames **E** and **F** where tumor cells surround clusters of normal cells. Finally, at the steady state regime the spatial domain shall be mostly occupied by tumor cells with the presence of few normal cells and empty vertices. It is also important to note that in a given tissue there are no empty spaces between cells, and in our simulations empty vertices are a consequence of the chosen exclusion diameters.

**Figure 3.**
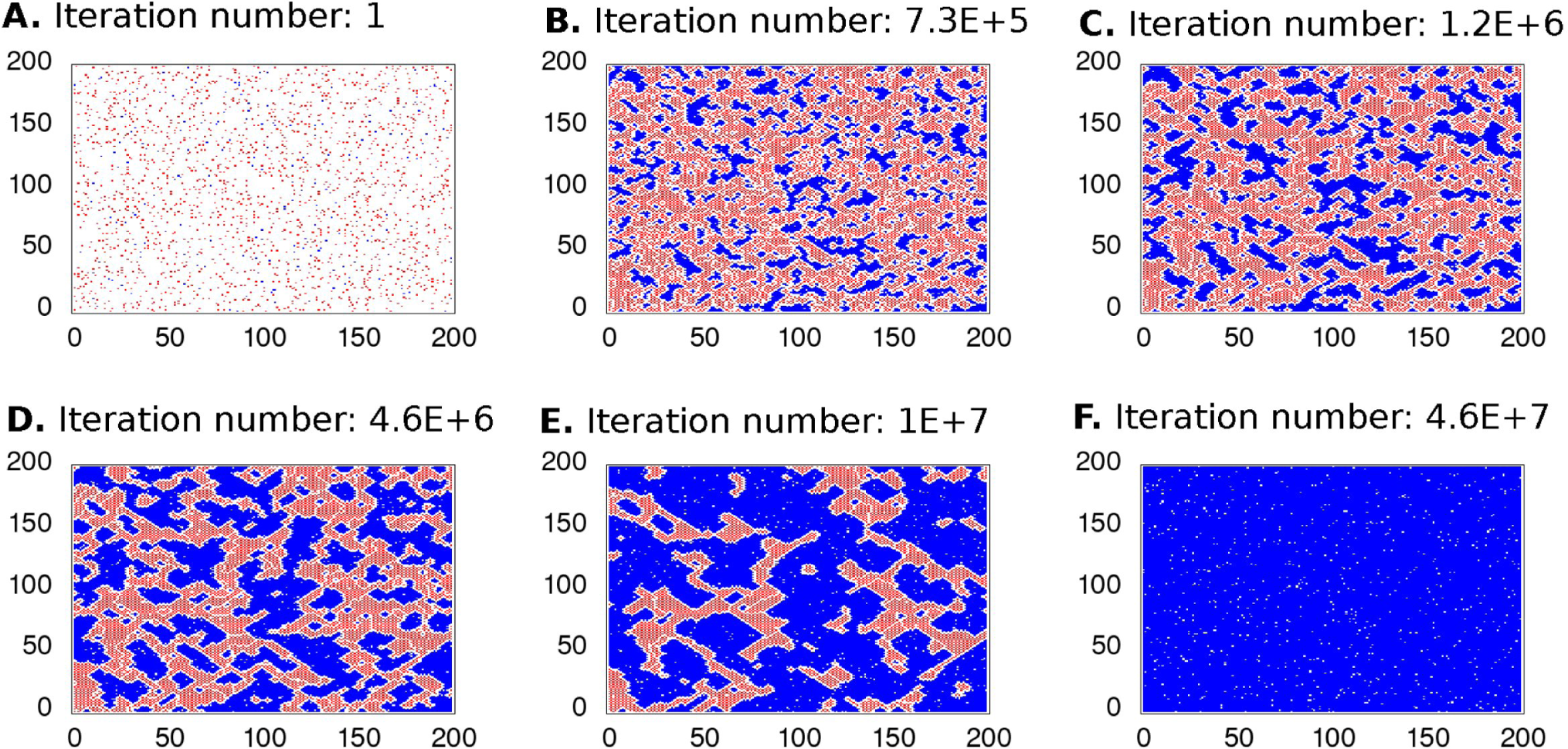
Cell occupation dynamics of the stochastic model. We present the spatial pattern dynamics with the same parameters of simulations as in Fig. 2. Simulation was conducted until a steady state has been achieved. The initial configuration of the spatial domain is presented in the left superior frame while the right inferior frame indicates the steady state configuration. The frames **A**–**D** corresponding to Fig. 2 while frames **E**, **F** show configurations after 10^5^ time steps of the Markov chain. The tumor cell population becames greater than that of the normal cells until they occupy most of the spatial domain.

The pictures on frames **E** and **F**, do not conform with the co-culture experiments presented here. Their presence has two purposes, the first is to show an asymptotic behavior of our model and how one might expect a co-culture experiment to behave when the only difference between the tumor and normal cells was their degree of *allelophilia*. A disagreement between experimental results and simulations can be explained by the fact that the experiments have been conducted for relatively short time-periods. Also, the proposed stochastic model might miss some important aspects of the co-culture development (e.g., because the two cell types are different on more aspects than their degree of contact inhibition). See the *Discussion* section of the paper.

##### A robustness analysis of prevalence of the most tolerant cell type

Fig. 4 shows results of an analysis of prevalence of the most tolerant species after 3 billion iterations. Frame **A** indicates the average density at the final iteration after 100 repetitions on the *y* axis and multiple values for the ratio *α*_2_/*α*_1_, with error bars included for each value of the ratio. The average densities are approximately the same for the two cell types when *α*_2_/*α*_1_ ~ 6 and cell type two prevails for *α*_2_/*α*_1_ = 10 at the final iteration. Frame **B** comprises three samples of the Markov chain simulation displaying the cell densities when *α*_2_/*α*_1_ = 6. The three samples have a qualitatively similar behavior but each of them shows a difference in the degree of prevalence of the cell type 1 due to the randomness of simulations. The final spatial configurations for the three samples are shown in frames **C**, **D** and **E**. The different patterns can be explained by large error bars indicated on frame **A**. Frame **C** shows a coexistence-like pattern where type two cells surround cells of type one. Frame **D** displays a condition where type two cells still prevail while prevalence of type one cells is observed in frame **E**. Results in frames **B** and **E** indicate that for a larger amount of iterations one may expect the most tolerant cell type to prevail in a spatial domain; increasing the ratio *α*_2_/*α*_1_ causes this phenomenon to take longer to happen. Frame **F** presents an analysis of melanoma cell clusters in the experiments and simulations. The top graphs show that the cluster areas in both simulations and experiments have similar box-plots. Here 1 pixel^2^ of area of the heatmap obtained from the simulation is equated with ~ 93 *μ*m^2^ of the area in the experiment. The square root is used in the identification of the perimeter correspondence. Notice that there is a good agreement between the median of both aspect ratios and convex hull obtained from both experiments and simulations (See the *Statistical Analysis* section at the Supplementary Information). Furthermore, frame **F** shows that clusters have a diversity of shapes and sizes, randomly distributed over the plate and the simulation grid. This emphasizes stochasticity of the observed phenomenon.

**Figure 4.**
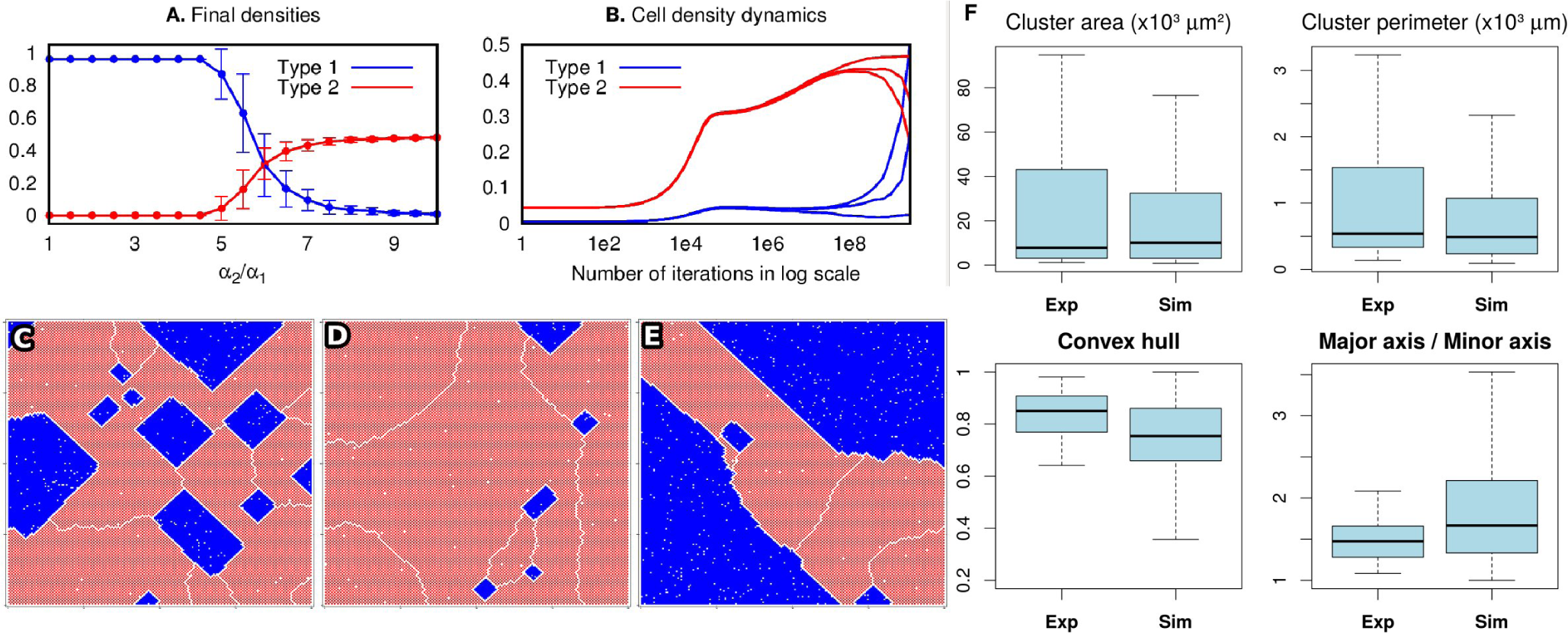
Analysis of the impact of variation of the value *α* upon the dynamics of cell proliferation. **A**. The density of the two cell types after 3·10^10^ iterations for *α*_1_ = 0.1 and the ratio *α*_2_/*α*_1_ varying from 1 to 10. The remaining parameters of the model assume the same values as in Figs 2 and 3 and the densities are obtained for 100 repetitions of the simulations. **B**. Three terminal configurations obtained in three simulation rounds: they exhibit similar qualitative behavior. However, differences in the sequence of pseudo-random numbers result in different times to achieve a prevalence of one cell type over the other. **C**,**D**,**E**. A spatial configuration of the cells at the final iteration corresponding to the three patterns of **B**. Note separating white lines (empty layers) between parts of the normal (red) cell population: these lines indicate boundaries of ‘phases’ (even and odd checker-boards). This is in agreement with the statements of Theorem 1 from the supplementary information. **F**. The analysis of cluster properties as obtained from the experimental data and simulations. We have obtained a good agreement between the box-plots for the cluster area, perimeter, convex hull and aspect ratio (p > 0.01) when we take 1 pixel on a simulation heatmap to be ~ 10 *μ*m.

#### The distance distribution quantifies the degree of *allelophilia* of the cells

To evaluate the degree of tolerance between cells of the same type, we performed an image-based analysis of the distribution of the distances between keratinocytes or melanoma cell lines in the co-culture. From the distribution of distances between cells we defined a minimal distance allowed for cells of the same type as the most frequent distance observed (a mode of the histogram). The distance distribution of keratinocytes assumes only distances appearing within the same range as those found in melanoma cell clusters. Three different distance distribution signatures have been observed (see Fig. 5). (a) A pre-confluence state where no significant difference has been detected between cells due to cell spread and low density (Fig. 5**A**). (b) At confluence, melanoma clusters are formed, and it is possible to detect a difference between histograms of distance distributions (Fig. 5**B**). (c) In a post-confluence state, a peak in the histogram for melanoma cells has been observed (Fig. 5**C**), showing a shorter distance more frequent than for keratinocytes. Tumors are denser than the surrounding normal tissue. This indicates a high degree of proximity and therefore a greater tolerance between cancer cells.

**Figure 5.**
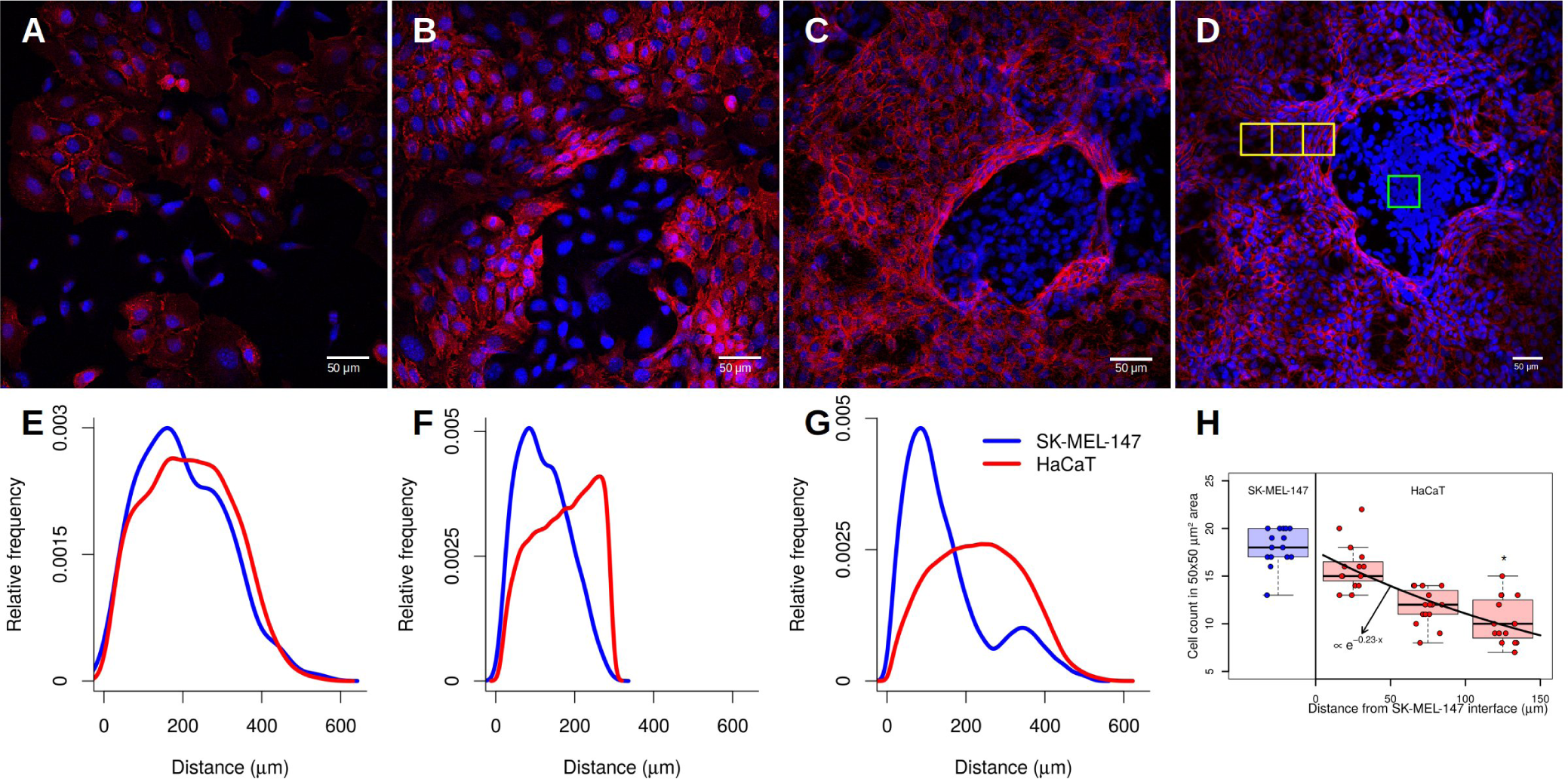
Representative histogram of the cell-to-cell distances distribution. **A**. A pre-confluent state of HaCaT and SK-MEL-147 cells in co-culture. **B**. At confluence, cells occupy all space available and the SK-MEL-147 cell cluster can be observed. **C**. A post-confluence state shows a high density of SK-MEL-147 cells with HaCaT surrounding. **D**. The distribution of the distances of the cells in the non-confluent state shows no difference between the cell types. **E**. In the confluent state, cell distance distribution begins to show a difference in the more frequent distance between cells of the same type. **F**. SK-MEL-147 cells tend to aggregate and form clusters surrounded by HaCaT cells. The distance distribution shows a higher relative frequency of shorter distances of SK-MEL-147 than HaCaT cells. Red and blue lines represent the distribution of HaCaT and SK-MEL-147 cell lines respectively. **G**. Representative image of an SK-MEL-147 cell cluster surrounded by HaCaT cells. Green squares indicate a high density of SK-MEL-147 cells and yellow squares indicate HaCaT cells areas with different densities at distances of 0-50, 51-100 and 101-150 *μ*m from SK-MEL-147 cells. **H**. The keratinocytes density dependence on the distance from the interface of the melanoma clusters. Boxplots indicates the median and interquartile range for SK-MEL-147 (blue) and HaCaT (red) cell count in a 50×50 *μ*m^2^ area from 15 selected locations of duplicated experiments. The data were fitted via an exponential decay using the one-way ANOVA and Tukey’s post-test between HaCaT and SK-MEL-147 densities (p < 0.001)

#### Keratinocyte density depends on the distance from melanoma

To evaluate melanoma proliferation restriction we quantified the density of keratinocytes in three distinct regions surrounding the tumor cells at 50, 100, and up to 150 *μ*m and at the high density cancer cells domain (Fig. 5**G**). As one can see, the keratinocyte density is affected by the distance from the melanoma cell domain (Fig. 5**H**). This morphology of epithelial cells are observed when cells are at high contact of inhibition. The keratinocytes are affected by proximity to melanoma cells, and more distant cells as a transitional morphology towards more dispersed state. Such a pattern is repeated and is induced by the communication between the SK-MEL-147 and HaCaT cells. A change on the proliferation rates in co-culture experiments indicates that the interaction between the tumor and normal cells is affecting their growth capabilities. As our data show, cells rearrange their spatial patterns to ensure that an initially well-mixed population of two cell types turns into clusters of a single cell type, either normal or tumor.

## Discussion

### The novelty of this work is that quantitative co-culture experiments are combined with a specific mathematical model

The co-culture experiments demonstrate that the low sensitivity to contact inhibition is responsible for a decrease in proportion rates of keratinocytes to melanoma cells and formation of tumor-like melanoma clusters. The explanation of these features has been provided through a specific mathematical model.

Our aim was to understand the role of a low degree of contact inhibition on the development of cancer *in situ*. The melanocyte and keratinocyte interaction through a direct contact is an important mechanism controlling their proliferation^27^. We observed a decrease on the proportion of keratinocytes to melanoma in the course of time: see Fig. 1**C**. Furthermore, a development of spatial pattern was observed, progressing from a well-mixed configuration (Fig. 5**A**) towards dense melanoma clusters surrounded by spatially spread domains with a relatively high density of keratinocytes (Fig. 5**C**).

To explain the observed phenomena, we use a mathematical model with a particular form of interaction between cells of the same and different types through different degrees of contact inhibition. The melanoma cells are supposed to have a lower level of contact inhibition than the keratinocyte cells. A mathematical mechanism of a stochastic nature has been considered, based on a Widom-Rowlinson-type model and its modifications^24–26.^ The model has its own subtleties as stressed in the supplementary information. For the chosen minimal distances *D*(1, 1) = 1, *D*(2, 2) = 1, *D*(1, 2) = *D*(2, 1) = 3 the model explains all possible types of terminal configurations.

The dynamics is presented as a Markov chain on admissible cell configurations with specified exclusion diameters for the keratinocyte (healthy/normal) and melanoma (tumor) cells; in the simulations we use the same rates for supposed transitions for cells of both types: division, death, and migration. Numerical simulations have been performed, confirming a number of features observed in the co-culture experiments. Tumor cells have smaller exclusion diameters; according to our interpretation, they are less sensitive to contact inhibition than healthy ones.

A major question is about including mechanical aspects of the cell motions into the model. This approach have been employed in investigations of homeostasis and morphogenesis^28,29^ and has shaded light on how mechanic stress affects contact inhibition molecular mechanisms^30^. Such an implementation in our stochastic approach would need a serious re-vision of the model which could be a topic for future research. In the mechanical approach cell-to-cell interaction happens through ‘springs’ which compression (dilation) controls cells growth rates^31, 32^.Our approach corresponds to the limit for which the ‘spring’ is fully compressed and it generates an infinite potential barrier, that is, the exclusion diameters are establishing a hard core potential for cell-to-cell interaction.

The proposed theoretical approach enabled us to reproduce a temporal increase on the melanoma proportion in a series of numerical simulations. A formation of spatial patterns on the two-dimensional grid has been also confirmed, albeit not at a desired level of accuracy. An existing agreement between the co-culture experiments and numerical simulations suggests that the development of a co-culture of melanoma and keratinocyte cells has a distinctive stochastic aspect which can be quantified. This may lead to a new methodology of prognosis for the development of carcinoma *in situ*, particularly formation of tumor-type clusters of melanoma cells and a collective reaction of keratinocyte cells to such a formation. Specific biochemical mechanisms of cellular interaction standing behind these phenomena need further studies. This would involve a variety of mathematical models modifying the present model. In particular, exclusion areas for healthy cells may vary and depend upon their distances from developing clusters of cancer cells.

Our characterization of the degree of contact inhibition suggests the use of the cells-to-cells distances to quantify sensitivity to contact inhibition. Tissues with tumors are denser than the normal ones^33–35^ and, thus, tumor cells are expected to be closer to each other forming geometrically packed arrangements. This is confirmed in Fig. 5**F** where we evaluate the distance distributions for keratynocytes and melanoma cells after confluence. The distribution of melanoma cell-to-cell distances is displaced to the left in comparison with the keratinocytes; this reinforces our interpretation of the exclusion diameters as a measure of the degree of contact inhibition of a cell.

The present experiments have been conducted with co-culture cell mono-layer. In the mathematical model it was interpreted as a stochastic dynamics of a two-dimensional admissible configuration on a grid. The passage to dimension three is mathematically possible (at a cost of an increase in the simulation time). We expect the qualitative features of the model presented here to be preserved. Such an extension would enable us to evaluate experiments on the formation of *in vivo* carcinoma *in situ* and to compare experimental results with simulations, aiming to establish a quantitative model to investigate carcinogenesis at its early stage. However, it would require new methods of marking and counting cells in the co-culture.

### Contact inhibition and the necessity for its full characterization

The theoretical curve for the ratio between the tumor and normal cell density has a more pronounced shape in comparison with the experimental data (Figs. 3**B** and 1**C**). This may be attributed to other effects controlling cell proliferation in the co-culture. A more precise correspondence between experimental and theoretical results might be established by engineering two cell lines with very similar proliferation, death, and migration rates but having different degrees of modulation of contact inhibition. This could have practical applications on the design of cancer treatment aiming to increase the degree of contact inhibition of tumor cells ^4, 36, 37^.

The simulation based on a mathematical model of a realistic cell interaction is quite promising: the higher level of *allelophilia* of the tumor cells may be a key feature for their prevalence within tissues. This can be detected through exclusion diameters as measure of the degree of contact inhibition of cell proliferation. Other types of mutual influence could also be taken into account.

The co-culture experiments show that the mechanism of a higher allelophilic degree might be due to molecular complexes surrounding the cells (see Fig. 5G and 5H). This shows the necessity of establishing a biochemical characterization of the exclusion diameters (or more factors) involved in the cell interaction.

### A view in the general context of cancer research

Let us address some aspects of possible criticism of our approach. We recognise that an excessive use of analogies between biological phenomena taking place at different scales (e.g., tissue and ecological levels) is risky. On the other hand, the evolutionary paradigm is an established tool for understanding carcinogenesis, provided that a proper caution is taken ^18,21,38^. In particular, while comparing ecological phenomena with those at a tissue level one must consider the specificity of cell-to-cell interactions and propose proper forms for quantitative modeling. An important mechanism here is the presence of membrane molecules causing the less immunogenic cancer cells to be selected for survival^39,40^. Another example is the development of cancer cells capable of performing glycolysis in hypoxic environment, when tumors reach a critical size interpreted as an adaptation ^41,42^. Such a new feature of cancer cells causes local acidosis toxic to normal cells; this is considered as an analog of allelopathic interactions between plants or bacteriocins in microtube^43^. In contrast, the present study focuses on an impact of an allelophilic behavior upon the formation of a carcinoma *in situ* which, within the evolutionary paradigm, indicates that the selective advantage of cancer cells may happen because of their greater degree of allelophilia.

## Methods

### Cell culture

The Supplementary information file contains details on cell culture proliferation assay. Human immortalized keratynocites (HaCaT) and human metastatic melanoma (SK-MEL-147) cell lines were used in this study. Cells were grown in Dulbecco’s Modified Eagle’s Media (DMEM) supplemented with 10% fetal bovine serum (FBS) in tissue culture incubator (5% CO2 at 37°C).

### CDH1 immunofluorescence

HacaT and SK-MEL-147 cells were seeded over 24-well plate with coverslips and grown for eight days. Cells were fixed with ice-cold methanol: acetone (1:1) at 4°C for 15 min. Cells were blocked with 5% bovine serum albumin (BSA) in phosphate-buffered saline for 1 h at room temperature. After blocking, cells were incubated with antibody against CDH1 (1:200 in 1 % BSA, Transduction Laboratories) for 45 min at room temperature. After washing, the coverslips were then incubated with Alexa Fluor 546 secondary antibody (1:400, Invitrogen) and 50 *μ*g/mL Hoechst 33258 (Invitrogen) in 1% BSA for 2 h at room temperature. Immunofluorescence images for each day of experiment (*n*=30) were acquired with an Nikon Eclipse E600 microscope, under 20x objective. The cells were counted based on the number of nuclei present in an image and the distance between cells were measured based on the centroid point of the image of the cell nucleus. The nuclei count were made manually using Cell Counter plug-in in ImageJ software (See the *Image Analysis* section in supplementary information file for more details).

### Cluster features

To evaluate melanoma cell clusters features, we performed an analysis of 60 clusters at the 8th day of the experiment. Shape descriptions performed on images captured under 10x objective consist of area, perimeter, convex hull 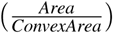 and aspect ratio (*Major axis /Minor axis*). Features of the clusters of cells of type 1 presented by one hundred repetitions of the simulations were also measured in the same way in heatmaps of 10^5^ iterations.

### Data availability statement

The data generated during and/or analysed during the current study are available from the corresponding author on reasonable request.

## Acknowledgements

This study was supported by CAPES (Process n. 88881.062174/2014-01, AFR). IS thanks the Math Department, University of Denver, for support and hospitality. AUS thanks Programa de Educaço Tutorial – MEC/SESu for support. YS thanks the Math Department, Penn State University, for support and attention. We thank Prof. Marsha Rosner for careful reading and suggestions that improved the quality of this manuscript. The authors are thankful for anonimous reviewers whose comments enhanced manuscript’s quality.

## Author contributions statement

(1) All authors contributed to the formulation of the cancer cell proliferation model as presented in the paper; (2) IS and YS were responsible for the initial mathematical model of tumor generation including the Markov chain algorithm; (3) the Markov chain algorithm was modified and used for simulations by AFR, AUS, WWL, ASQ, MCCM, RC; (4) MCCM, TCT, RC, AFR, AUS, ASQ, WWL were responsible for the biological experiments, data collection and initial interpretation; (5) All authors discussed the results and their final interpretation; (6) All authors participated in writing and revising the manuscript.

## Additional information

### Supplementary Information

regarding the results are available at http://www.nature.com/srep

## References

1. Abercrombie, M. Contact inhibition and malignancy. Nat. 281, 259–262 (1979).

2. Hanahan, D. & Weinberg, R. A. Hallmarks of cancer: the next generation. Cell 144, 646–674 (2011).

3. Stockinger, A., Eger, A., Wolf, J., Beug, H. & Foisner, R. E-cadherin regulates cell growth by modulating proliferation-dependent beta-catenin transcriptional activity. The J. Cell Biol. 154, 1185–1196 (2001).

4. Seluanov, A. et al. Hypersensitivity to contact inhibition provides a clue to cancer resistance of naked mole-rat. Proc. Natl. Acad. Sci. United States Am. 106, 19352–19357 (2009).

5. Gatenby, R. A. & Maini, P. K. Mathematical oncology: Cancer summed up. Nat. 421, 321–321 (2003).

6. Anderson, A. R. A. & Quaranta, V. Integrative mathematical oncology. Nat. Rev. Cancer 8, 227–234 (2008).

7. Kreeger, P. K. & Lauffenburger, D. A. Cancer systems biology: a network modeling perspective. Carcinog. 31, 2–8 (2010).

8. Byrne, H. M. Dissecting cancer through mathematics: from the cell to the animal model. Nat. Rev. Cancer 10, 221–230 (2010).

9. Kuang, Y., Nagy, J. D. & Eikenberry, S. E. Introduction to Matematical Oncology (CRC Press, 2015), 1 edn.

10. Bozic, I., Allen, B. & Nowak, M. A. Dynamics of targeted cancer therapy. Trends molecular medicine 18, 311–316 (2012).

11. Tomasetti, C. & Vogelstein, B. Variation in cancer risk among tissues can be explained by the number of stem cell divisions. Sci. 347, 78–81 (2015).

12. Patel, A. A., Gawlinski, E. T., Lemieux, S. K. & Gatenby, R. A. A cellular automaton model of early tumor growth and invasion. J. Theor. Biol. 213, 315–331 (2001).

13. Anderson, A. R. A., Weaver, A. M., Cummings, P. T. & Quaranta, V. Tumor morphology and phenotypic evolution driven by selective pressure from the microenvironment. Cell 127, 905–915 (2006).

14. Gatenby, R. A. & Gawlinski, E. T. A reaction-diffusion model of cancer invasion. Cancer Res. 56, 5745–5753 (1996).

15. Rejniak, K. A. & Anderson, A. R. A. Hybrid models of tumor growth. Wiley Interdiscip. Rev. Syst. Biol. Medicine 3, 115–125 (2011).

16. Hatzikirou, H., Basanta, D., Simon, M., Schaller, K. & Deutsch, A. ’Go or grow’: the key to the emergence of invasion in tumour progression? Math. medicine biology: a journal IMA 29, 49–65 (2012).

17. Gatenby, R. A. & Frieden, B. R. Application of information theory and extreme physical information to carcinogenesis. Cancer Res. 62, 3675–3684 (2002).

18. Gatenby, R. A. & Vincent, T. L. An evolutionary model of carcinogenesis. Cancer Res. 63, 6212–6220 (2003).

19. Gatenby, R. A. & Frieden, B. R. Inducing catastrophe in malignant growth. Math. Medicine Biol. 25, 267–283 (2008).

20. Gatenby, R. A., Silva, A. S., Gillies, R. J. & Frieden, B. R. Adaptive Therapy. Cancer Res. 69, 4894–4903 (2009).

21. Arnal, A. et al. Evolutionary perspective of cancer: myth, metaphors, and reality. Evol. Appl. 8, 541–544 (2015).

22. Delbrück, M. Statistical fluctuations in autocatalytic reactions. J. Chem. Phys. 8, 120–124 (1940).

23. Puliafito, A. et al. Collective and single cell behavior in epithelial contact inhibition. Proc. Natl. Acad. Sci. USA 109, 739–744 (2012).

24. Widom, B. & Rowlinson, J. New model for the study of liquid-vapor phase transition. J. Chem. Phys. 52, 1670–1684 (1970).

25. Mazel, A., Suhov, Y., Stuhl, I. & Zohren, S. Dominance of most tolerant species in multi-type lattice Widom-Rowlinson models. J. Stat. Mech. Theory Exp. 2014, P08010 (2014). ArXiv: 1403.5825.

26. Mazel, A., Suhov, Y. & Stuhl, I. A Classical WR Model with q Particle Types. J. Stat. Phys. 159, 1040–1086 (2015).

27. Cichorek, M., Wachulska, M., Stasiewicz, A. & Tyminska, A. Skin melanocytes: biology and development. Adv. Dermatol. Allergol. Dermatol. I Alergologii 30, 30–41 (2013).

28. Guillot, C. & Lecuit, T. Mechanics of epithelial tissue homeostasis and morphogenesis. Sci. 340, 1185–1189 (2013).

29. Streichan, S., Hoerner, C., Schneidt, T., Holzer, D. & Hufnagel, L. Spatial constraints control cell proliferation in tissues. Proc. Natl. Acad. Sci. USA 111, 5586–5591 (2014).

30. Pan, Y., Heemskerk, I., Ibar, C., Shraiman, B. & Irvine, K. D. Differential growth triggers mechanical feedback that elevates hippo signaling. Proc. Natl. Acad. Sci. USA 113, E6974–E6983 (2016).

31. Shraiman, B. I. Mechanical feedback as a possible regulator of tissue growth. Proc. Natl. Acad. Sci. USA 102, 3318–3323 (2005).

32. Kim, N.-G., Koh, E., Chen, X. & Gumbiner, B. M. E-cadherin mediates contact inhibition of proliferation through hippo signaling-pathway components. Proc. Natl. Acad. Sci. USA 108, 11930–11935 (2011).

33. Eisenhoffer, G. T. & Rosenblatt, J. Bringing balance by force: live cell extrusion controls epithelial cell numbers. Trends Cell Biol. 23, 185–192 (2013).

34. Butcher, D. T., Alliston, T. & Weaver, V. M. A tense situation: forcing tumour progression. Nat. Rev. Cancer 9, 108–122 (2009).

35. Kumar, S. & Weaver, V. M. Mechanics, malignancy, and metastasis: the force journey of a tumor cell. Cancer Metastasis Rev. 28, 113–127 (2009).

36. Zeng, Q. & Hong, W. The emerging role of the hippo pathway in cell contact inhibition, organ size control, and cancer development in mammals. Cancer Cell 13, 188–192 (2008).

37. Harvey, K. F., Zhang, X. & Thomas, D. M. The Hippo pathway and human cancer. Nat. Rev. Cancer 13, 246–257 (2013).

38. Breivik, J. The evolutionary origin of genetic instability in cancer development. Semin. Cancer Biol. 15, 51–60 (2005).

39. Zhang, K., Lu, Q., Zhang, Q. & Hu, X. Regulation of activities of {NK} cells and {CD4} expression in t cells by human hnp-1, -2, and -3. Biochem. Biophys. Res. Commun. 323, 437–444 (2004).

40. Dunn, G. P., Old, L. J. & Schreiber, R. D. The three es of cancer immunoediting. Annu. Rev. Immunol. 22, 329–360 (2004).

41. Gatenby, R. A. & Gillies, R. J. Why do cancers have high aerobic glycolysis? Nat. Rev. Cancer 4, 891–899 (2004).

42. Lash, G. E. et al. Oxygen as a regulator of cellular phenotypes in pregnancy and cancer. Can. J. Physiol. Pharmacol. 80, 103–109 (2002).

43. Crespi, B. & Summers, K. Evolutionary biology of cancer. Trends Ecol. & Evol. 20, 545–552 (2005).

